# A Deep Learning Model for Predicting Tumor Suppressor Genes and Oncogenes from PDB Structure

**DOI:** 10.1101/177378

**Authors:** Amirhossein Tavanaei, Nishanth Anandanadarajah, Anthony Maida, Rasiah Loganantharaj

## Abstract

While cancer is a heterogeneous complex of distinct diseases, the common underlying mechanism for uncontrolled tumor growth is due to mutations in proto-oncogenes and the loss of the regulatory function of tumor suppression genes. In this paper we propose a novel deep learning model for predicting tumor suppression genes (TSGs) and proto-oncogenes (OGs) from their Protein Data Bank (PDB) three dimensional structures. Specifically, we develop a convolutional neural network (CNN) to classify the feature map sets extracted from the tertiary protein structures. Each feature map set represents particular biological features associated with the atomic coordinates appearing on the outer surface of protein’s three dimensional structure. The experimental results on the collected dataset for classifying TSGs and OGs demonstrate promising performance with 82.57% accuracy and 0.89 area under ROC curve. The initial success of the proposed model warrants further study to develop a comprehensive model to identify the cancer driver genes or events using the principle cancer genes (TSG and OG).

## 1 Introduction

Common themes (https://www.cancer.gov/about-cancer) among many different types of cancer at the molecular level include (1) mutations in proto-oncogenes that alter the function of the regular cell cycle to uncontrollable cell division, (2) mutations in cancer suppressor genes that alter their cell regulatory mechanism, and (3) mutations in DNA-repair genes that cause further mutations in cells instead of repairing them. Traditional machine learning algorithms such as decision trees, random forests (RF), artificial neural networks (ANN), support vector machines (SVM) have been successfully applied to build predictive models for various aspects related to cancer including prognosis of cancer, classification of cancer types from data sources such as clinical data, SNP’s, gene expressions [1, 2, 3, 4, 5]. Recently, deep learning [6, 7] has shown remarkable performances for predicting the specificity of DNA and mRNA binding sites [8], functional classification [9], protein folding pattern [10], and for cancer categorization [11, 12]. Automatic detection and prediction of the either oncogenes or cancer suppression genes from their three dimensional features is a big step in discovering their structural characteristics to improve the state-of-the-art in making a dent in cancer treatments. To our knowledge there is no or few works done in applying machine learning in identifying oncogenes or cancer suppression genes from the three dimensional structures.

Although there are many different cancer types such that finding a coherent pattern representing their drivers is challenging, cancer manifests as tissue grows in an uncontrolled manner due to malfunction in the regular cell cycle process. Along with many other factors, it has been documented through experimentation that mutations in proto-oncogenes and in tumor suppressor genes and their regulatory mechanism play major roles in tumor growth. OGs refer to the genes that increase the number of cells while TSGs refer to the genes that control the cell growth process. Osborne et al [13] reviewed common OG and TSG malfunctions in human breast cancer. The OG/TSG detection improves the cancer identification performance as discussed in [14]. They have used genomic data and their variants from the cancer genomic atlas (TCGA), ICGC and COSMIC and have applied a random forest model integrating five statistical tests to detect cancer genes and specify them as likely OG and TSG. A question arises that how to classify OGs and TSGs only from their three-dimensional protein structures and the biochemical properties of the amino acids that forms the structure without extra statistical tests or other feature extraction modules? Prediction of the functional annotation of proteins is being improved by various methods such as prediction by sequence similarity [15, 16], evolutionary relations [17], genetic interactions [18], protein-protein interactions [19], protein structures and gene-ontology hierarchy [9, 20, 21].

In this paper, we propose a deep convolutional neural network (CNN) to classify TSGs and OGs based on their PDB structures and the biochemical properties. CNNs have shown high performances in visual feature extraction and classification [22, 23]. Additionally, CNNs provide hierarchical feature extraction modules which are robust against rotation, scale, and local translation. Thus, these types of visual extraction modules can be used to discover discriminative information of the PDB structures by mapping the biochemical properties annotated with 3-D atomic coordinates appearing on the outer surfaces to visual feature maps.

## 2 TSG and OG Dataset Preparation

### 2.1 Protein Structure

The tertiary protein structure is determined by three-dimensional geometric shape with a single polypeptide backbone. It contains a variety of bonding interactions between the side chains on the amino acids. Figure 1 shows a protein structure in which the colors exhibit its secondary structure. In this paper, we concentrate on the protein’s atomic coordinates appearing on the outer surfaces (x,y,z) and their associated biochemical properties.

**Figure 1:**
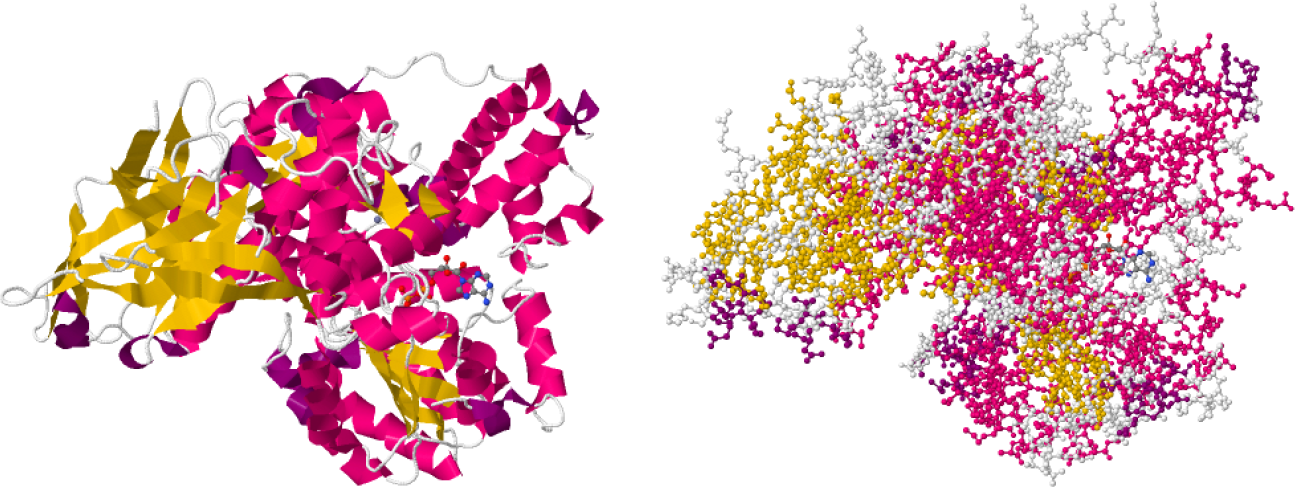
Tertiary structure of protein ‘4CDG’. Cartoon (left) and atomic (right) displays.

### 2.2 Biochemical Features

Experimentally annotated genes that are related to cancer are downloaded from COSMIC (https://cancer.sanger.ac.uk/census) V82 for human (GRCh38). For the purpose of this machine learning experiment in identifying the genes role in cancer from their 3D structure, we have focused on tumor suppressor genes and oncogenes. The recent version of the COSMIC annotated gene lists has 137 TSG and 78 oncogenes. These gene sets are combined together and are clustered with DAVID Bioinformatics Resources 6.8 [24] using only direct annotation from gene ontology (http://www.geneontology.org) and other functional categories provided by the tool. The gene ontology [25] provides a structured vocabulary to annotate genes and their products by providing three orthogonal ontologies: biological process (BP), cellular components (CC), and molecular function (MF), each of which is modeled as a directed acyclic graph. As expected, the TSGs and the OGs are clustered into non-overlapping, separate clusters. This confirms that they are appropriate candidates for separate functional predictive models.

The Ensembl ids of OGs and TSGs are mapped to the PDB ids by using the UniProt web tool [26] (http://www.uniprot.org/uploadlists). The PDB files were downloaded from the protein data bank website (https://www.rcsb.org/pdb/download/download.do#Structures). The PDB format contains a standard format for macromolecular structure data obtained by X-ray diffraction and NMR studies [27].

To interpret and distinguish these genes, we provide a feature extraction module to represent morphological characteristics of their tertiary structure. The feature extraction module has two folds: 1) identifying the surface *C*_*α*_ atoms; 2) extracting the surface atoms’ properties. The pseudo code of feature extraction is shown in Figure 2.

**Figure 2:**
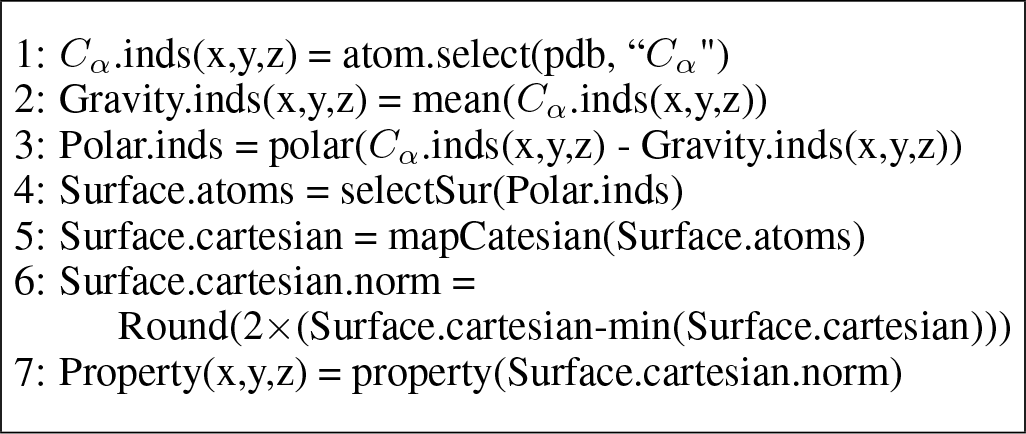
Pseudo-code for PDB feature extraction.

#### 2.2.1 Surface *C*_*α*_ Identifying

For each PDB file, the surface *C*_*α*_ atoms are chosen. To find the surface atoms, the Cartesian coordinates are changed to polar coordinates from the centroid and then, with 1 degree resolution, the highest radius atom is selected as the surface atom (Figure 2 lines 1 through 5). Finally, the surface indices < *x*, *y*, *z* > are converted to decimal numbers starting from < 0, 0, 0 > (Figure 2 line 6).

#### 2.2.2 Atom Properties

The PDB files provide information for amino-acids placed in the *C*_*α*_ coordinates. In our model, each PDB file is represented by sixteen features along with their 3-D coordinates. These 16 features are extracted according to the corresponding amino-acids collected from http://www.proteinstructures.com/Structure/Structure/amino-acids.html. The sixteen features that are contributed to the 20 amino-acids are shown in Table 1. These features are used in the “property” function as shown in Figure 2: line 7. *Note:* the feature values of Pka-NH2 and P-Ka-COOH are normalized in the range [0, 1].

**Table 1.**
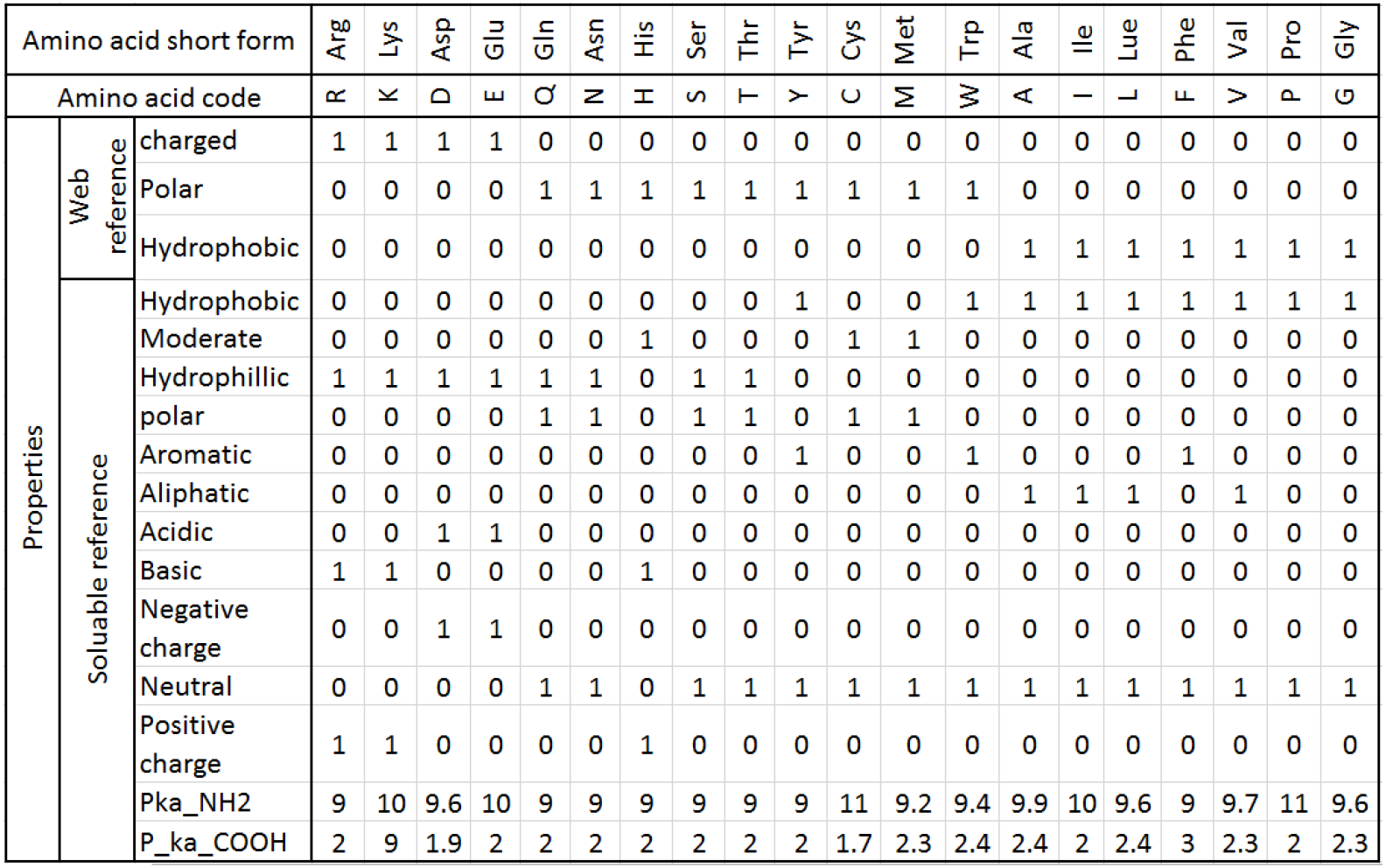
Amino-acids and their associated properties.

## 3 Deep Learning Model

In this section, we first explain the data processing steps required for preparing the feature maps fed into the CNN; and then, the network architecture is explained.

### 3.1 Input Feature Maps

As mentioned earlier, each PDB file is represented by 16 features associated with the atomic coordinates < *x*,*y*,*z* >. To covert the three dimensional feature space to the feature maps (2-D), we generate three independent feature sets associated with three atomic projections on < *x*, *y* >,< *y*, *z* >, and < *x*, *z* > feature spaces. Therefore, each PDB file can be converted to three perpendicular 2-D feature spaces. In the next step, each projection is converted to 16 feature maps corresponding to the sixteen feature values computed in the previous section.

This approach converts a 3-D structure to three feature map sets with dimensions of 200 × 200 × 16 pixels (16 feature maps of 200 × 200). Processing the projections is much faster than processing the 3-D structures while not losing information considerably due to the PDB’s sparse structure. Furthermore, each feature map set of a projection denotes specific features of the protein while preserving its spatial information. Figure 3 shows the feature maps of one of the three projections (< *x*, *y* >) for a TSG.

**Figure 3:**
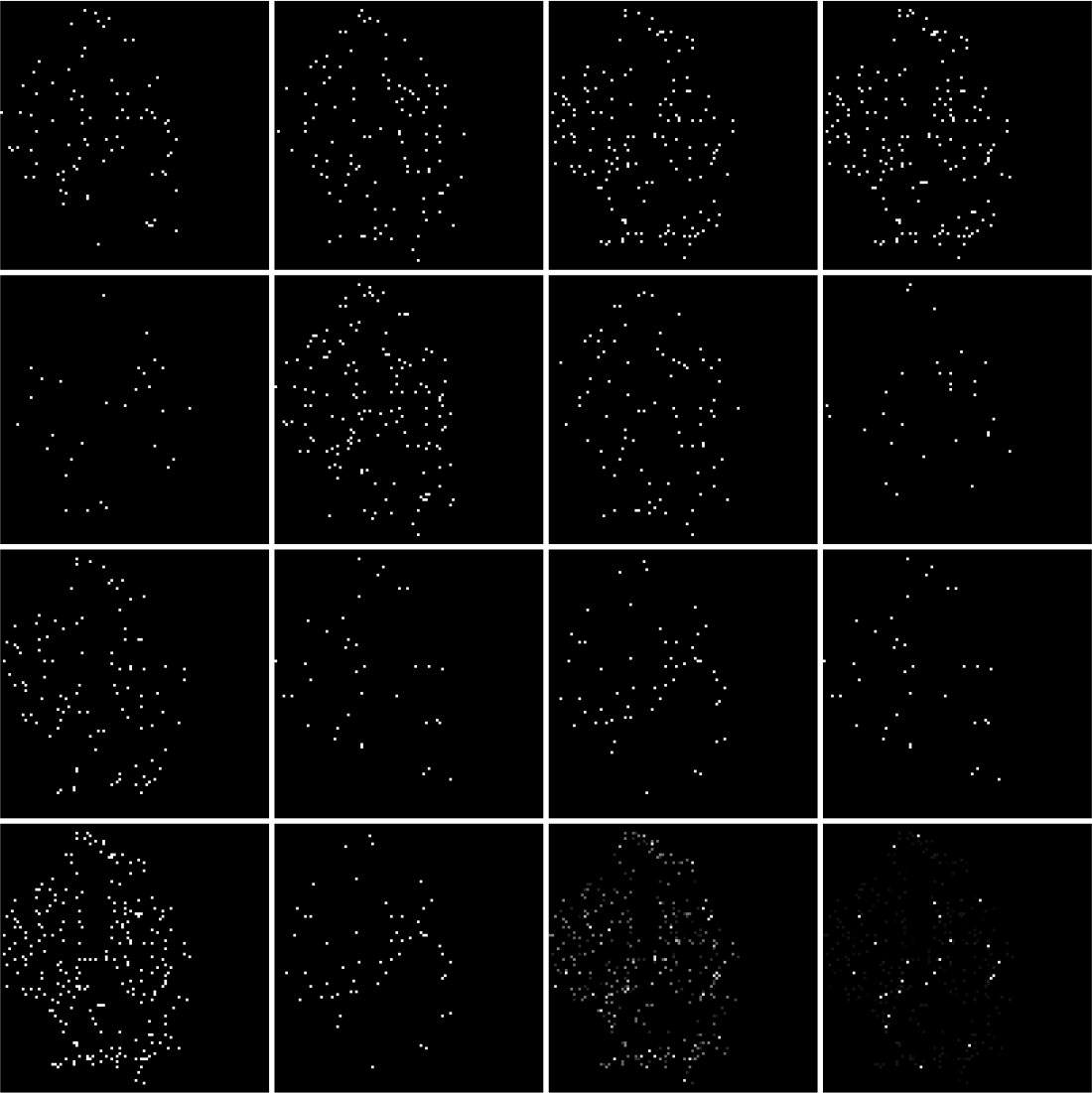
Sixteen feature maps obtained from the < *x*, *y* > projection of protein ‘4CDG’ (a TSG). Each map illustrates one of the biological features calculated in the previous section. The row-wise map order is analogous to the order of features shown in Fig. 4.

### 3.2 CNN Architecture

The deep learning model, in this study, develops a parallel CNN with three branches followed by a multi-layer fully connected neural network. Figure 4 shows this deep CNN’s architecture. The model consists of four convolution and pooling layers and three fully connected layers including the final classifier. The convolution kernel size (*p*), pooling strides (*s*_*i*_), number of hidden neurons (*h*_1_, *h*_2_), convolution padding (*γ*), and the number of generated feature maps (*d*_*i*_) are shown in Table 2. These parameters have been set up after a number of control experiments and initial evaluations.

**Figure 4:**
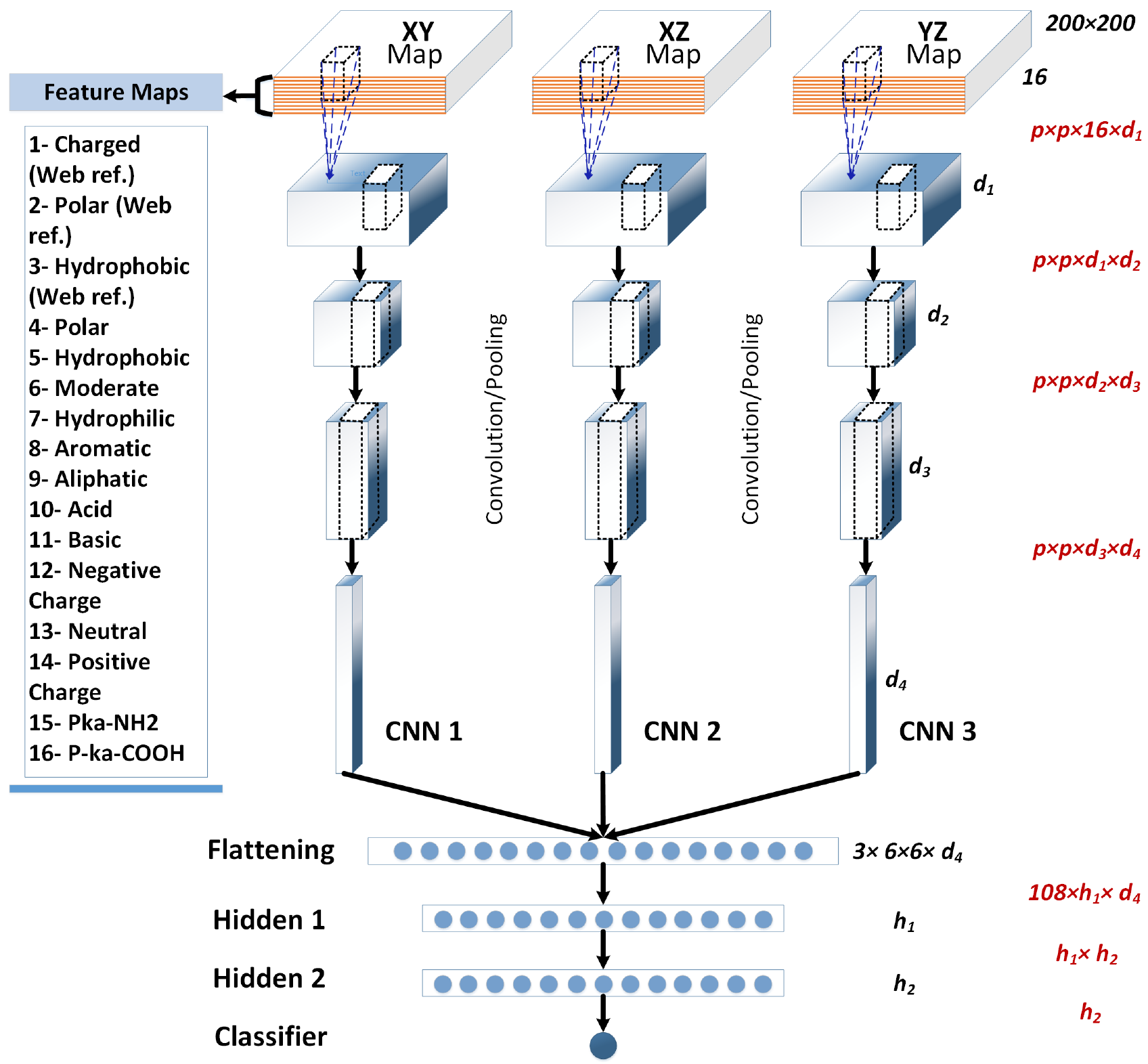
The deep CNN’s architecture for OG/TSG prediction. For simplicity, the convolution and pooling layers are shown as one module. The red expressions calculate the number of trainable parameters for each layer.

**Table 2.** The CNN’s parameters.

| Parameter | Value | Parameter | Value |
| --- | --- | --- | --- |
| $d_1$ | 32 | $s_1$ | 4 |
| $d_2$ | 32 | $s_2$ | 2 |
| $d_3$ | 64 | $s_3$ | 2 |
| $d_4$ | 64 | $s_4$ | 2 |
| $p$ | 3-9 | $\gamma$ | 2 |
| $h_1$ | 100 | $h_2$ | 50 |

As shown in Figure 4, the CNN receives three 200 × 200 × 16 feature maps in parallel and performs a binary (TSG/OG) classification. Each layer is equipped by the rectified linear unit (ReLU) activation function. We used 30% dropout in the fully connected layers to control probable over training. More details about the network’s training process is discussed in the next section (Experiments). The features utilized for generating the feature maps are shown on the left side of the network (Figure 4). The convolution/pooling layers extract 108 × 64 = 6912 visual features. The number of trainable parameters are shown on the right side of the network. If we consider *p* = 5, we will have 888, 250 trainable parameters.

## 4 Experiments and Results

The proposed model is evaluated on the dataset that we collected in Section 2. The dataset consists of 2379 PDB files (1191 TSG and 1188 OG) that is converted to 7137 feature maps with 16 channels. The 2379 feature map sets, each representing one particular protein structure, are randomly divided into separate training and testing sets with 2029 training and 350 testing samples. This dataset division method is repeated three times using different random seeds. Finally, the model is trained and evaluated over 100 iterations.

We implemented the model using the Torch library [28] available via: https://github.com/tavanaei/Cancer-Suppressor-Gene-Deep-Learning.

### 4.1 Results

We ran four evaluations on the CNNs with different convolution kernel sizes (*p* = {3, 5, 7, 9}) to find a proper patch size for extracting the feature maps’ visual features. Figure 5 illustrates the CNNs’ accuracy rates over 100 training iterations. It is shown that the 3 × 3 patch is not able to discover the protein maps’ features well. The best accuracy belongs to the networks with convolution kernels with *p* = {7, 9} patch sizes. Table 3 shows the detailed performance measures of the proposed model. The best performed model reported accuracy rate of 82.57% and area under the ROC curve (AUROC) of 0.89. The test sets used to generate Figure 5 and Table 3 are slightly different (fewer samples were used for the plot).

**Figure 5:**
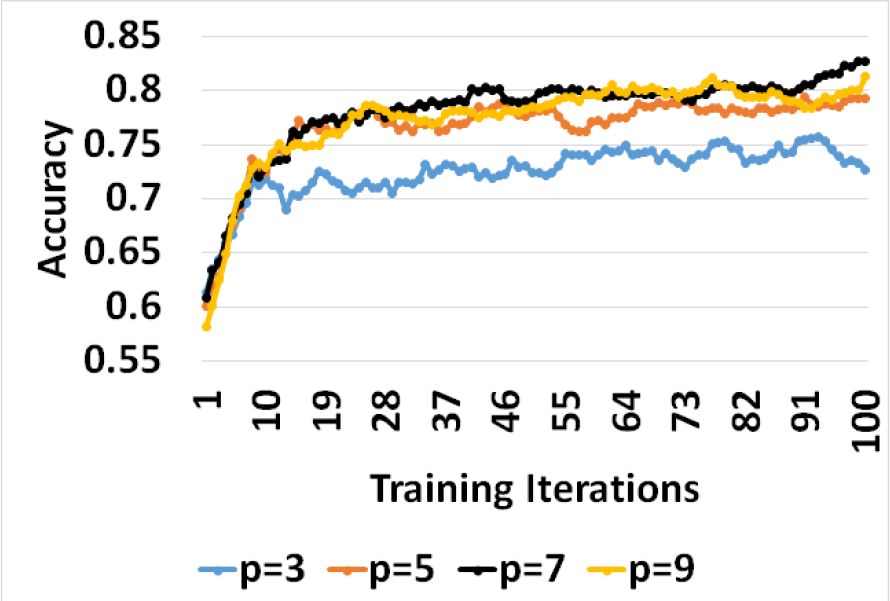
Accuracy of the networks with *p* = {3, 5, 7, 9} convolution patch sizes over training. *μ* = 0.05

**Table 3.** Performance of the CNNs in terms of accuracy, precision, recall, and area under ROC (AUROC). *μ* = 0.05.

| $p$ | 5 | 7 | 9 |
| --- | --- | --- | --- |
| **Accuracy (%)** | 81.71 | **82.57** | 79.43 |
| **Recall (%)** | 85.80 | 81.87 | 80.70 |
| **Precision (%)** | 78.38 | 82.35 | 77.97 |
| **AUROC** | 0.881 | **0.887** | 0.869 |

To asses the model’s convergence speed, Figure 6 shows the model’s performance with respect to different learning rates. The models trained by the learning rates *μ* > 0.03 reach accuracy rates higher than 75% after 20 iterations. The best performing models were trained using learning rates of 0.05 and 0.06. Table 4 also shows high performances for these learning rates while it is evaluated on a slightly different test set (same as Table 3).

**Figure 6:**
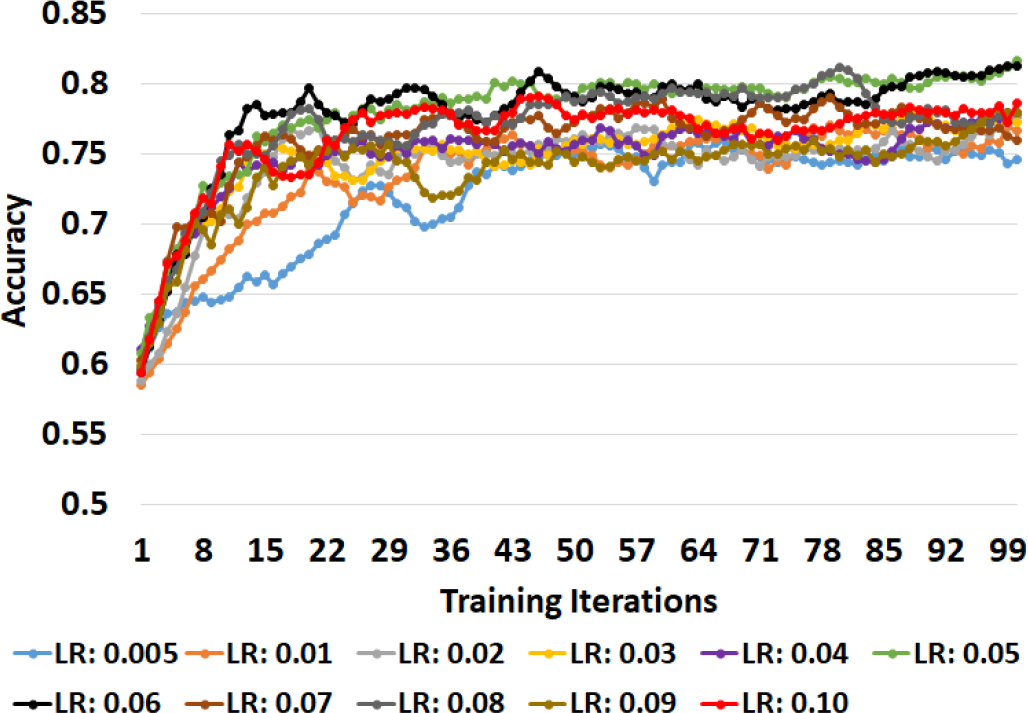
Accuracy of the CNN over training versus learning rate. *μ* = 0.05 and *p* = 7.

**Table 4:** Performance of the CNNs versus learning rate. Convolution patch size *p* = 7.

| Learning Rate | Accuracy | Recall | Precision | AUROC |
| --- | --- | --- | --- | --- |
| **0.005** | 78.29 | 86.98 | 73.13 | 0.880 |
| **0.010** | 77.71 | 81.66 | 74.59 | 0.871 |
| **0.020** | 80.00 | 85.80 | 75.92 | 0.876 |
| **0.030** | 79.14 | 83.43 | 75.81 | 0.878 |
| **0.040** | 80.57 | 80.00 | 80.00 | 0.872 |
| **0.050** | **82.57** | 81.87 | 82.35 | **0.887** |
| **0.060** | 82.00 | 85.12 | 79.01 | **0.883** |
| **0.070** | 79.14 | 79.29 | 77.91 | 0.877 |
| **0.080** | 82.00 | 82.84 | 80.46 | 0.880 |
| **0.090** | 78.57 | 85.21 | 74.23 | 0.871 |
| **0.100** | 80.00 | 81.66 | 77.97 | 0.874 |

### 4.2 Summary and Discussion

The raw input data are obtained from the 3D structures of proto-oncogenes and cancer suppression genes. The function and the biological processes of a protein are dependent on the outer surface configuration and their biochemical properties. We have developed an algorithm to identify atoms located on the outer surface and have used selected sixteen properties of the amino acids as shown in Table 1. The 3D configuration of the outer surface is mapped onto three orthogonal planes. Each property becomes a channel in the feature map of the CNN as illustrated in Figure 4. The proposed CNN model with the 16 channel feature map achieved a remarkable performance of 82.6% accuracy rate and 0.89 AUROC. The model becomes very useful in annotating uncharacterized PDB structures into either the TSG or the OG structures.

Furthermore, this performance of our model compares favorably with the statistical methods studied by [14] on pan-cancer genome sequencing data [29] which consists of very rich genomic information. Event though the dataset we used for evaluating our model is different from their dataset. Table 5 compares the AUROC value reported in our study with the AUROC values reported by state-of-the-art statistical methods for OG versus TSG identification. Our model outperforms the six out of eight methods and is close to the best AUROC.

**Table 5:** AUROC of OG/TSG identification using the statistical methods reported by [14] and our model.

| Method | Description | AUROC |
| --- | --- | --- |
| Truncation Rate | Rate of truncating events* | **0.922** |
| Unaffected Residues | Intra-gene mutation clustering/recurrence* | 0.479 |
| VEST Mean | Functional impact bias* | 0.71 |
| Patient Distribution | Bias in patient labels* | 0.556 |
| 'Cancer Type Distribution | Cancer type bias* | 0.612 |
| Oncodrive-fm | Gonzalez-Perez and Lopez-Bigas [30] | 0.725 |
| Oncodrive-CLUST | Tamborero et al., [31] | 0.597 |
| Random Forest | Ensemble on the first 5 methods (*) | **0.924** |
| Our Model | DCNN applied to the PDB structure | **0.887** |

## 5 Conclusion

A deep learning approach was proposed in this paper to classify the cancer genes: proto-oncogenes and tumor suppressor genes. By having a model that confidently identifies proto-oncogene or cancer suppressor genes from the structure and the biochemical properties of amino acids, we are providing a general methodology that will lead to a new tool to discover a new set of cancer suppressor genes or proto-oncogenes that may not have been identified in the literature of having such functionality. By activating such dormant cancer suppression gene through drugs, we improve the chances of controlling tumor growth. Of course, the identified potential cancer suppression genes have to be verified through testing with rat or mouse model which resemble human gene content.

This investigation was established in two steps: 1) Protein feature extraction from the PDB tertiary structure; 2) modeling the gene patterns using a parallel deep convolutional neural network (CNN). The proposed DCNN preserves the spatial information of the tertiary structure while modeling the protein structure/features via three parallel, independent visual feature extraction modules. Finally, the fully connected neural network of the DCNN classifies the combined visual features.

The experimental results showed high performance of 82.57% and 0.887 accuracy rate and area under the ROC curve, respectively. The initial success of our model warrants our future study to apply the same deep learning approach on new datasets for predicting different cancer types to identify the cancer drivers.

